# CryoEM map of *Pseudomonas aeruginosa* PilQ enables structural characterization of TsaP

**DOI:** 10.1101/2020.05.29.123786

**Authors:** Matthew McCallum, Stephanie Tammam, John L. Rubinstein, Lori L. Burrows, P. Lynne Howell

**Author notes:** These authors contributed equally to this work (Matthew McCallum, Stephanie Tammam). Department of Biochemistry, University of Washington, Seattle, WA 98195- 7350, USA. Lead contact: PLH.

## Abstract

The type IV pilus machinery is a multi-protein complex that polymerizes and depolymerizes a pilus fibre used for attachment, twitching motility, phage adsorption, natural competence, protein secretion, and surface-sensing. An outer membrane secretin pore is required for passage of the pilus fibre out of the cell. Herein, the structure of the tetradecameric secretin, PilQ, from the *Pseudomonas aeruginosa* type IVa pilus system was determined to 4.3 Å and 4.4 Å resolution in the presence and absence of C_7_ symmetric spokes, respectively. The heptameric spokes were found to be the two tandem C-terminal domains of TsaP. TsaP forms a belt around PilQ and while the protein is not essential for twitching motility, over-expression of TsaP triggers a signal cascade upstream of PilY1 leading to cyclic di-GMP up-regulation. These results resolve the identity of the spokes identified with Proteobacterial PilQ homologs and may reveal a new component of the surface-sensing cyclic di-GMP signal cascade.

**IMPACT STATEMENT:** The type IV pilus is critical for bacterial virulence. The co-structure of the pilus secretin PilQ and TsaP is determined. Characterization of TsaP implicates it in surface-sensing signal transduction.

## INTRODUCTION

The type IVa pilus (T4aP) facilitates many bacterial functions including attachment, twitching motility, phage adsorption, natural competence, protein secretion, and surface-sensing^1,2^. The surface-sensing role of T4aP is of particular interest due to its involvement in the up-regulation of cyclic di-GMP – the master regulator of biofilm formation^3,4^. Biofilms are of critical medical and industrial importance, as they increase bacterial tolerance of otherwise uninhabitable conditions, antibiotics, and to the immune system^5^.

The T4aP machinery comprises a cell envelope-spanning multi-protein complex. This machinery polymerizes long thin fibres that attach to surfaces and retract to pull bacteria towards the point of attachment^1,2^. The pilus is a polymer mainly composed of the major subunit, PilA. At the pilus base, the inner membrane protein PilC and cytoplasmic ATPase motors, PilB and PilT, catalyze polymerization and depolymerization, respectively^6,7^. Minor pilins and PilY1 initiate pilus polymerization and are thus thought to localize at the pilus tip^8,9^. The pilus exits the periplasm through an outer membrane pore formed by PilQ oligomers. The PilQ pore is aligned with the motors by a subcomplex of PilM, PilN, PilO, and PilP^10,11^. PilF shuttles PilQ to the outer membrane and facilitates its oligomerization. PilMNOP, PilY1, and PilT are all required for T4aP-mediated surface sensing ^12,13^. It is unclear whether other proteins that are part of the T4aP apparatus and interface with PilM, PilN, PilO, and PilP – such as PilQ – are also needed for surface sensing.

PilQ and PilF are the secretin and pilotin of the T4aP, respectively. Secretins and their cognate pilotins create outer-membrane channels in various secretion and pilus systems. Secretin oligomers have a unique double-layered β-barrel architecture^14,15^. The outer β-barrel forms a large vestibule that passes though the outer membrane^15^. The inner β-barrel curves inwards to form a gate at the midway point of the secretin β-barrel^15^. Domains N1 and N0 form rings below the β-barrel^15^. PilQ secretins from Proteobacteria, structurally characterized at low resolution, show radial spokes emerging from a central barrel^16-19^. Models of PilQ from *Thermus thermophilus* (PilQ^Tt^) and *Vibrio cholerae* (PilQ^Vc^) competence pilus systems – which are divergent from Proteobacterial T4aP secretins – were recently built into 7 and 2.7 Å resolution electron cryomicroscopy (cryoEM) maps, respectively ^20,21^. These competence secretins lack the lip subdomain and the radial spokes found in T4aP secretins.

Structures of the related type II secretion system (T2SS) secretins also have radial spokes^22,23^. These structures were determined at 3-4 Å resolution, permitting modeling of the spoke protein as the T2SS pilotin^22,23^. This finding suggests that the spokes in the T4aP structures could be the T4aP pilotin, PilF. However, PilF is structurally unrelated to the T2SS pilotin, and other studies indicate that the spokes are most likely the protein TsaP^16,19^. 2D class averages of cryoEM particles embedded in purified *Neisseria* sp. membranes show spokes around the PilQ barrel, and these spokes disappear upon deletion of TsaP^16^. Similarly, electron cryotomography of *Myxococcus xanthus* show that density consistent with the location of the spokes around PilQ disappears upon deletion of TsaP^19^. Little is known about the molecular function of TsaP, except that it has a LysM domain characteristic of septal peptidoglycan binding^16^. Neither TsaP nor PilF are not expressed in the same operon as PilQ.

We previously resolved a 7.4 Å map of PilQ from *Pseudomonas aeruginosa* (PilQ^Pa^) with presumed tetradecameric stoichiometry and *C*_7_ symmetric spoke density^18^. Although this was the highest resolution structure of a secretin at that time, it was not sufficient for model building. Herein, our reprocessing of those micrographs with improved image analysis and model-building algorithms yielded three distinct structures of PilQ^Pa^. These maps were of sufficient resolution to enable building of an atomic model. Mass spectrometry revealed the identity of the protein bound in the spoke density to be TsaP, and a TsaP model was readily built into the spoke density. While probing for the function of TsaP, we found that that its overexpression triggers cyclic di-GMP up-regulation upstream of PilY1, implicating a role in the T4aP-dependent surface-sensing signal cascade.

## RESULTS

### PilQ^Pa^ forms tetradecameric and pentadecameric barrels

We previously resolved a 7.4 Å resolution PilQ^Pa^ cryoEM map^18^. Using the template-based automatic particle selection algorithm implemented in cryoSPARC v.2^24^, eight times more particles were selected automatically from the original micrographs than when particle selection was performed manually (**Supplementary Table 1**). *Ab initio* classification followed by heterogeneous particle refinement in 3D revealed three unique classes, with 44 % of particle images classified as *C*_14_ symmetric barrels with *C*_7_ symmetric radial spokes (**Figure 1A)**, 48 % of particle images classifying as *C*_14_ symmetric barrels without spokes (**Figure 1B)**, and 8 % as *C*_15_ symmetric barrels without spokes (**Figure 1C**). Refining the three classes yielded maps at 4.3, 4.4, and 6.9 Å resolution, respectively. The distinct *C*_14_ and *C*_15_ symmetric maps identified are reminiscent of the finding that the T2SS secretin can have more than one symmetry^25^.

**Figure 1.**
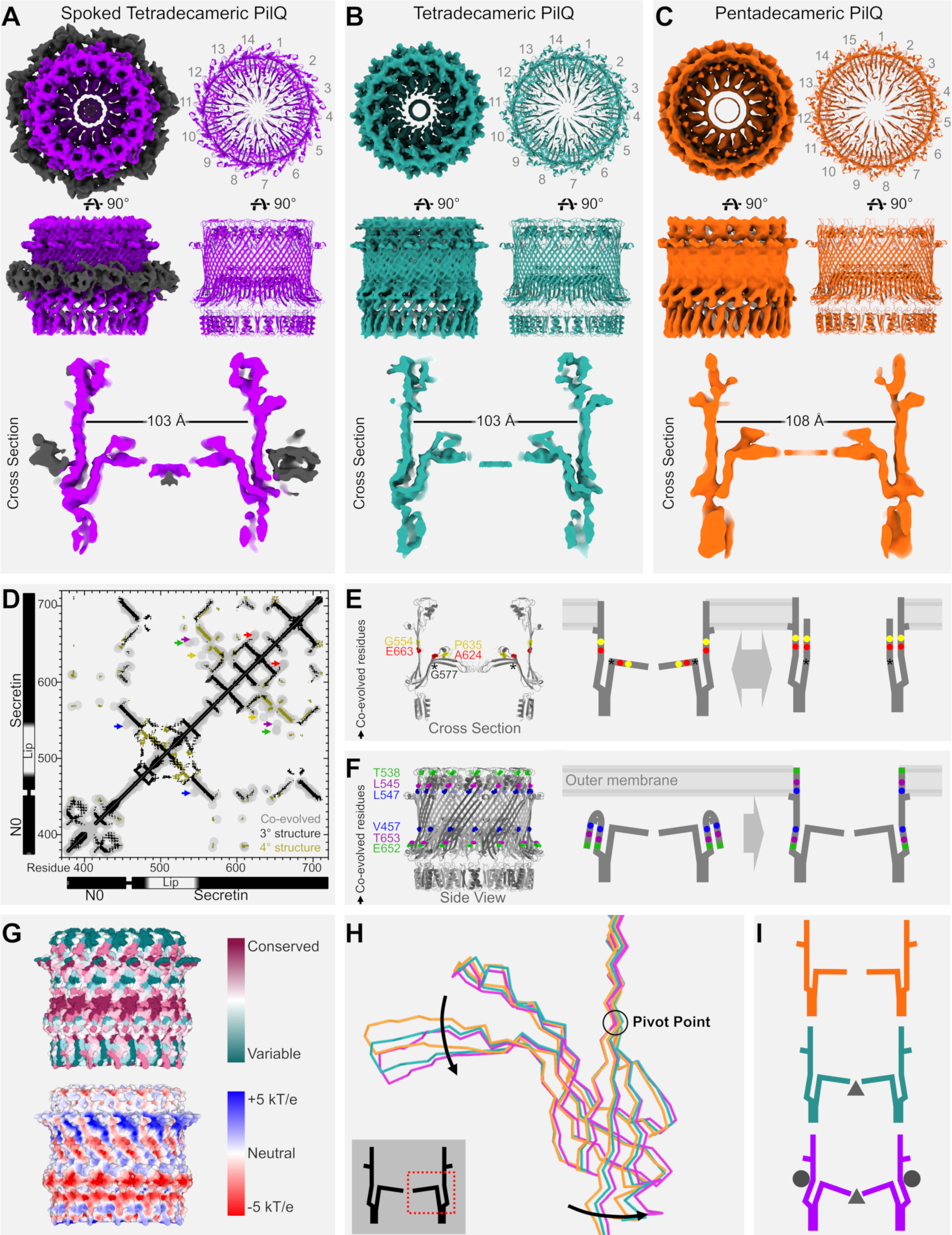
Electron cryomicroscopy of PilQ from *P. aeruginosa* reveals multiple conformers. **A-C**, Map (upper left) and model (upper right) of PilQ^Pa^ (coloured) with additional spoke and plug density (black). Top (top), side (middle), and cross section (bottom) views are shown. **A**, Map and model of spoked tetradecameric PilQ^Pa^ (purple). **B**, Map and model of tetradecameric PilQ^Pa^ without spokes (teal). **C**, Map and model of pentadecameric PilQ^Pa^ (orange). **D**, Evolutionarily coupled residues in PilQ (gray) are compared with the structure of PilQ^Pa^. Residue pairs that are within 8 Å of each other in the tertiary structure of PilQ^Pa^ are shown (black). Residue pairs that are within 8 Å of each other between PilQ^Pa^ protomers are also shown (gold). Specific evolutionarily coupled residues that are not supported by the PilQ^Pa^ structure are noted with coloured arrows: G554-P635 (yellow), E663-A624 (red), T538-E652 (green), L545-T653 (magenta), and L546-V457 (blue). **E**, Location of the G554-P635 and E663-A624 residue pairs, as well as hinge residue G577 (asterisk), on a cross section of PilQ^Pa^ (left) and a model for secretin gate opening consistent with these co-evolved residues. **F**, Location of the T538-E652, L545-T653, and L546-V457 residue pairs on PilQ^Pa^ (left) and a model for secretin barrel folding consistent with these co-evolved residues. **G**, Surface representation of conserved residues (top) and electrostatic potential (bottom) in PilQ^Pa^. **H**, Structure alignment of monomers from the spoked tetradecameric PilQ^Pa^ (purple), tetradecameric PilQ^Pa^ with no spokes (teal), and pentadecameric PilQ^Pa^ (orange) models. Grey square with black PilQ cartoon has a dotted red box highlighting area of PilQ shown. Structural differences are highlighted (arrows) and the pivot point for these differences is noted (black circle). **I**, Cartoon model summarizing the differences between the three PilQ^Pa^ structures determined. Pentadecameric PilQ^Pa^ (orange) is wider and has more horizontal secretin gates compared to the other structures. Tetradecameric PilQ^Pa^ (teal) has invaginated secretin gates and partial plug density (black triangle). Spoked tetradecameric PilQ^Pa^ (purple) has further invaginated secretin gates, plug density (black triangle), and spokes (black circles).

Local resolution in the barrel of the 4.3 and 4.4 Å maps is estimated to be 3.5 Å. The secondary structure and large side chains were visible in the map and enabled model building of 14 PilQ^Pa^ N1 and secretin domains (**Figure 1A**). Although we verified that the N0 and peptidoglycan-binding AMI_N domains of PilQ were present using mass spectrometry analyses (**Supplementary Figure 1**), they could not be modeled, consistent with these domains adopting random orientations relative to the secretin barrel in the absence of the peptidoglycan and the rest of the T4aP apparatus. Protomers of the barrel structure from the spoked barrel map were fit into the maps of the *C*_14_ and *C*_15_ barrels without spokes and refined.

The final models were compared to co-evolved residues within PilQ^Pa^ homologs as identified by the EVcouplings webserver^26^. Residues in proximity to one another in a structure develop complementary mutations as the protein evolves^27^. These co-evolved residues can be used to independently predict and validate the conformations of proteins^27^. We found a clear correlation between the co-evolved residues and the N1 and secretin domains, validating these structures (**Figure 1D**). The sequence of the lip subdomain is not well conserved in PilQ, and the poor sequence alignment in this region yielded fewer co-evolved residues in the lip subdomain (**Figure 1D**).

### Co-evolved residues are consistent with secretin gate and barrel dynamics

Co-evolved residues not directly supported by the model of PilQ^Pa^ were also identified, suggesting dynamics not visualized in the cryoEM maps. There are co-evolved residues between the secretin gate and the secretin barrel (**Figure 1E**), consistent with a swinging gate-like model of secretin gate opening demonstrated recently for the secretin from the type III secretion injectisome, InvG^28^. Residue G577 is located at the proposed hinge of the *P. aeruginosa* PilQ secretin gate (**Figure 1E**). Mutation of the corresponding residue in *M. xanthus* PilQ (G764S) causes vancomycin sensitivity^29^ presumably because the hinge of the gate is jammed open permitting entry of the bulky antibiotic across the outer membrane. This hypothesis is supported by the structure of a mutant of the T2SS secretin where the corresponding residue G458 was changed to alanine (G458A)^25^. The G458A T2SS secretin structure reveals a partially open secretin gate^25^.

Co-evolved residues were also identified at the top of the secretin barrel and near the bottom of the secretin barrel, adjacent to the secretin gate (**Figure 1F**). These co-evolved residues may be consistent with the secretin barrel folding such that the top of the secretin barrel covers the gate-adjacent segment of the secretin. The top of the secretin is hydrophobic (**Figure 1G**), like other secretin structures^15^, consistent with this region being the site for outer membrane insertion. Thus, one explanation for these co-evolved residues may be that a folding intermediate positions the top of the secretin barrel to mask these hydrophobic sites prior to membrane insertion (**Figure 1F**). The gate-adjacent section of the secretin barrel is highly conserved (**Figure 1G**) and provides the interface for the spoke density. Once PilQ inserts into the outer membrane and this region becomes exposed, the spoke protein could bind and stabilize the PilQ oligomer.

### Dynamics in the secretin gates visualized

The conformations of the secretin gates in the *C*_15_, *C*_14_, and spoked *C*_14_ PilQ^Pa^ barrels are different. While the secretin gate in the *C*_15_ barrels is perpendicular to the walls of the barrel, the secretin gate in the *C*_14_ barrel invaginates towards the periplasm. Comparing the *C*_15_ and *C*_14_ barrels without spokes, the residues at the center of the secretin gate are displaced by 4 Å (**Figure 1B-C, Animation 1**). This invagination is likely the result of the 5 Å reduction in diameter of the tetradecameric barrel relative to the pentadecameric form, leaving less room for the gate to extend perpendicularly from the walls of the barrel.

Comparing the *C*_14_ and spoked *C*_14_ barrels reveals that the secretin gate invaginates by an additional 3 Å in the spoked *C*_14_ barrel (**Figure 1A-B**). A smaller barrel diameter cannot explain this movement, as the diameters of the *C*_14_ barrel and spoked *C*_14_ barrels are identical. Comparing the *C*_15_, *C*_14_, and spoked *C*_14_ barrels reveals a pivot point for gate invagination in the secretin barrel adjacent to the secretin gate and the spokes (**Figure 1H-I and Animation 1**). This finding is consistent with the molecules bound at the spokes promoting secretin gate invagination.

### TsaP co-purifies with PilQ

There are two features in the maps of PilQ^Pa^ that cannot be explained by models of PilQ^Pa^ alone. First are the *C*7 symmetric spokes. The spoke density appears to be comprised of 14 structurally similar β-roll protein structures that form a belt around PilQ. There are differences in every other β-roll that give the spokes *C*7 symmetry (**Figure 1A**). There is also extra density beneath the secretin gate that plugs the small opening in the center of both tetradecameric PilQ barrels – the density is more pronounced in the spoked tetradecameric barrel map (**Figure 1A-B**). The local quality of the map beneath the secretin gate is too poor to identify distinct structural elements. Curiously, there is no evidence of *C*_15_ barrels with spokes or extra density beneath the secretin gates, hinting at stoichiometry-dependent interactions.

To identify the protein(s) that constitute the extra densities, mass spectrometry was performed on the protein sample used for cryoEM grid preparation. Aside from PilQ^Pa^, the most abundant components are PrpL, PA5122, and TsaP (**Supplementary Table 2, Supplementary Figure 2**). TsaP is the protein most likely responsible for the spoke density as it has been shown to be critical for PilQ spoke visualization in *Neisseria* sp. and *M. xanthus*^16,19^. PrpL is an extracellular protease secreted by the T2SS^30,31^ and thus unlikely to constitute these densities. PA5122 is an uncharacterized protein with a secretion signal sequence.

**Figure 2.**
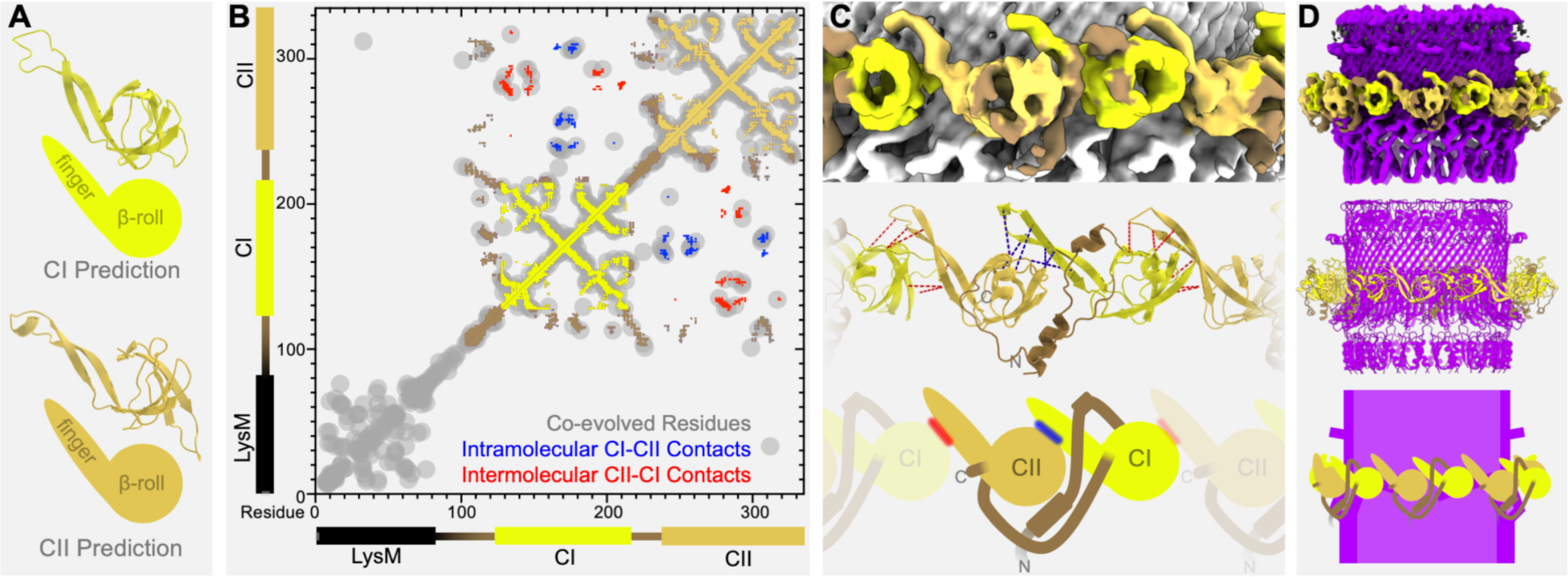
TsaP can be modeled in the PilQ spoke density as a heptameric ring. **A**, TsaP domain model predictions: the I-Tasser prediction of the CI domain (residues 126-211)^34^ and the Phyre^2^ prediction of the CII domain (residues 235-324)^33^. **B**, Evolutionarily coupled residues in TsaP (gray) are compared with the structure of the C-terminal 224 residues of TsaP. Residue pairs that are within 8 Å of each other are shown from the CI domain (yellow), the CII domain (wheat-yellow), and the linkers preceding and following the CI domain (brown). Intramolecular and intermolecular contacts in the TsaP structure between the CI and CII domains are also shown (blue and red, respectively). **C-D**, Map (top), model (middle), and cartoon (bottom) summarizing the structure of TsaP. Zoomed-in (**C**) and zoomed-out (**D**) views are shown. TsaP is coloured as in **B**; PilQ is shown in grey (**C**) or purple (**D**).

### Model building of TsaP reveals unique heptamer structure

The structure of TsaP is not known; however, TsaP was predicted to have an N-terminal peptidoglycan-binding LysM domain linked by unstructured residues to a C-terminal domain^19^. The C-terminal domain of TsaP, henceforth referred to as the CII domain, was predicted to be similar to the C-domain of the flagellar ring-forming component, FlgT^19^. The C-domain of FlgT has a β-roll structure with an extruding finger subdomain^32^ and the Phyre^2^ structure prediction of TsaP from *P. aeruginosa* is consistent with this fold (**Figure 2A**). To validate this result, TsaP co-evolved residues identified by the EVcouplings webserver^26^ were compared with the predicted structure of the TsaP CII domain (**Figure 2B**). The correlations between co-evolved residues and residues close to one another in the CII domain structure prediction provide support for its similarity to the FlgT C-domain. Inspection of TsaP co-evolved residues revealed a duplicate pattern of co-evolved residues preceding the CII domain (**Figure 2B**), consistent with a second domain with a similar structure, henceforth referred to as the CI domain. When the structure of the CI domain segment was predicted^33,34^ we found that it was also similar to the FlgT C-domain (**Figure 2A**). This is distinct from previous predictions that identified only a single C-domain in TsaP^19^.

The β-roll structure and extruding finger subdomain in the CI and CII models can be clearly seen in the spoke density (**Figure 2C**). There is also density consistent with linker residues preceding the CI domain and between the CI and CII domains. In contrast, Phyre^2^ predicted structures of PrpL and PA5122 do not fit the size or shape of the spokes. The C-terminal 224 residues of TsaP encompassing the CI and CII domains were therefore modeled and refined into the map. This model reveals that TsaP forms a heptameric ring with C7 symmetry around PilQ^Pa^ at the midpoint of the barrel where the internal gate is positioned (**Figure 2D**). Each finger subdomain contacts the adjacent C domain to mediate intramolecular CI^n^-CII^n^ interactions and intermolecular CII^n^-CI^n+1^ interactions. These finger subdomain interactions correlate with co-evolved residue pairs in TsaP (**Figure 2B**), further validating the model of TsaP built into this map.

Each C domain contacts a separate PilQ protomer. The buried surface area of the TsaP-PilQ interaction is 2924 Å^2^ per TsaP protomer, or 20468 Å^2^ for the entire 7:14 TsaP-PilQ interaction. The buried surface area of the TsaP-TsaP oligomeric interface is 913 Å^2^. The larger buried surface area of the TsaP-PilQ interface compared with the TsaP-TsaP oligomeric interface hints that the PilQ interaction may help facilitate TsaP oligomerization.

### Overexpression of TsaP triggers cyclic di-GMP signalling

*tsaP* mutants have a 40 % defect in twitching motility, which can be complemented *in trans* by leaky expression from *tsaP* under an arabinose-inducible promotor without the addition of arabinose (**Figure 3A**). We previously reported finding no obvious defect in twitching motility with the *tsaP* mutant^18^; this was likely the result of a limited number of replicates combined with the modest phenotype. Compared to the vector control, a 66 % defect in twitching motility is observed in wild-type *P. aeruginosa* or the *tsaP* mutant complemented with *tsaP* when induced with 0.1 % (w/v) arabinose – henceforth referred to as over-expressed TsaP (**Figure 3B**) – consistent with both deletion and over-expression of TsaP impairing twitching motility.

**Figure 3.**
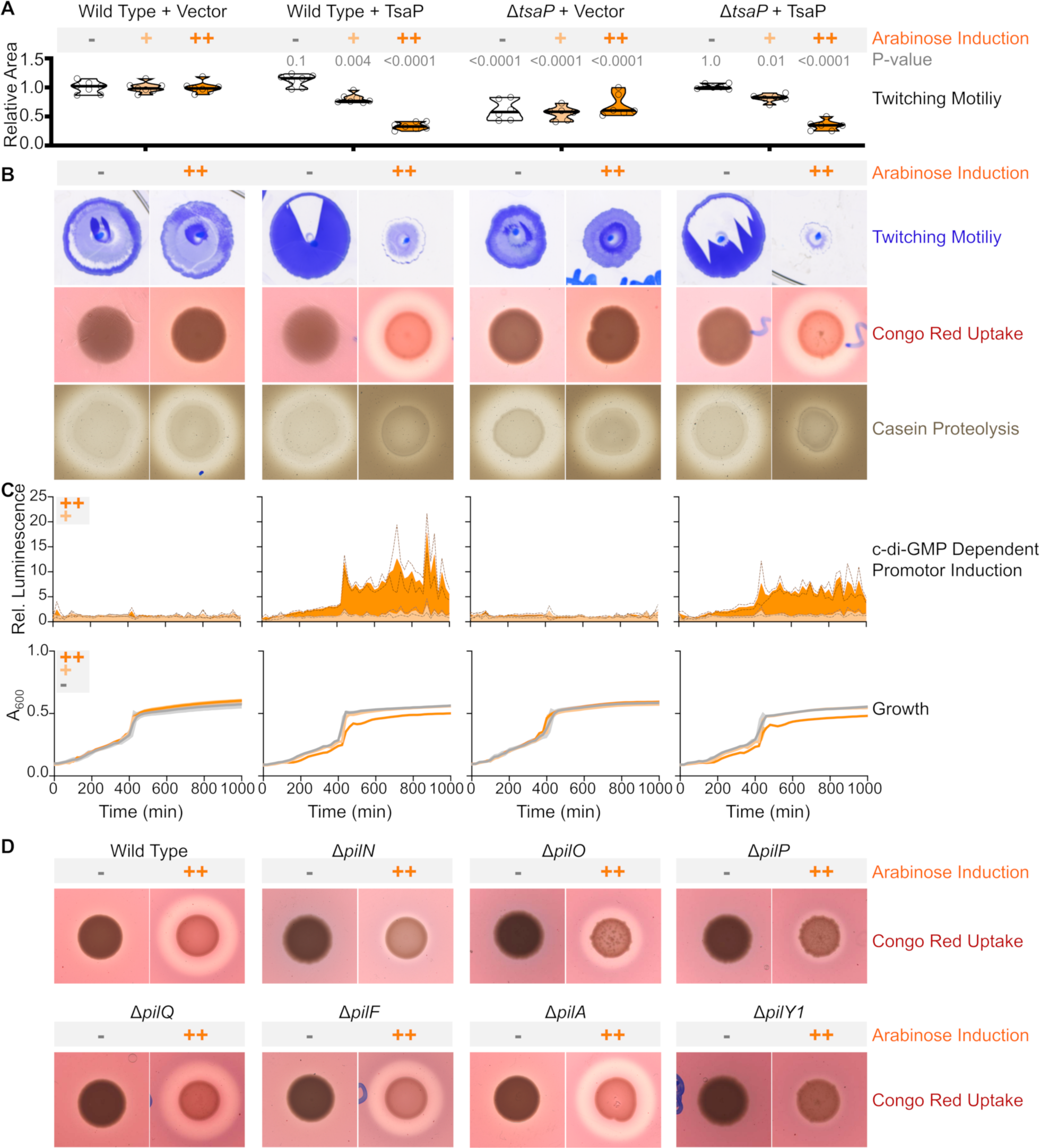
TsaP over-expression induces cyclic di-GMP up-regulation. Wild-type *P. aeruginosa* (left), the *tsaP* mutant (center left), *P. aeruginosa* expressing TsaP (center right), and the *tsaP* mutant expressing TsaP (right). **A**, Twitching motility was assayed by crystal violet staining and the areas traversed by the bacteria were measured (violin plot with individual measurements shown as empty circles; n = 7 for Wild-type with and without vector, n = 6 for the *tsaP* mutant with and without vector; data are representative of three independent trials). Mean twitching areas were compared to wild type plus the control vector at the corresponding arabinose concentration with the two-way ANOVA test; p-values are reported. **B**, Representative crystal violet stained twitching motility zones (top), Congo-red agar colonies (middle), and skim milk agar colonies (bottom) are shown for 0 % arabinose induction (-) or 0.1% arabinose induction (++). **C**, Luminescence over time relative to uninduced control without arabinose (top), with luciferase expression under the control of the cyclic di-GMP inducible *cdrA* promotor. Mean luminescence (n=3, representative of three independent trials) relative to a matched uninduced control is shown for 0.1% arabinose induction (orange, ++) or 0.01% arabinose induction (beige, +) as a filled in line with the standard error of the mean (dotted lines). Growth measured by absorbance at 600nm over time of liquid cultures (bottom). Mean growth (n=3, representative of three independent trials) is shown for 0 % arabinose induction (gray, -), 0.01% arabinose induction (beige, +), or 0.1 % arabinose induction (orange, ++) with the standard error of the mean shown (dotted lines). **D**, Congo red dye uptake assay of PAO1 (wild-type), PAO1::*pilN*, PAO1::*pilO*, PAO1::*pilP*, PAO1::*pilQ*, PAO1::*pilA*, PAO1::*pilF* and PAO1::*pilY1* with (++) and without (-) TsaP overexpression.

We also assessed the propensity of *tsaP* mutants and TsaP-overexpressing *P. aeruginosa* to form biofilms, using a Congo red dye uptake assay. Congo red binds to the Pel polysaccharide in *P. aeruginosa* biofilms, leading to red colonies and a small clear halo around the colonies where the dye has been taken up^35,36^. We found that the *tsaP* deletion mutant is indistinguishable from wild-type *P. aeruginosa* (**Figure 3B**). However, TsaP overexpression in wild type or the *tsaP* mutant leads to red colonies surrounded by pronounced clearing (**Figure 3B**), consistent with the over-expression of TsaP leading to increased biofilm formation.

To look at other surface-associated phenotypes, we assessed the propensity of TsaP overexpressing *P. aeruginosa* to secrete proteases using a casein proteolysis assay. Hydrolysis of casein by extracelluar proteases leads to clear halos around protease-secreting colonies on solid skim milk media^37^. Clear halos form around wild-type *P. aeruginosa* and the *tsaP* deletion mutant; however, no halos form around TsaP over-expressing colonies (**Figure 3B**), consistent with its over-expression decreasing extracellular protease secretion.

Reduced extracellular protease secretion, decreased twitching motility zones, and increased biofilm formation are all associated with cyclic di-GMP induction^4,38-40^. Measurement of luciferase expression from a cyclic di-GMP sensitive promotor verified that TsaP overexpression increases cellular cyclic di-GMP concentrations (**Figure 3C**). Growth curves confirmed that increased luciferase expression is not a result of increased cell growth (**Figure 3C**). These results are consistent with TsaP over-expression up-regulating the production of cyclic di-GMP.

### TsaP signalling occurs upstream of PilY1

We probed the dependency of cyclic di-GMP signal upregulation caused by TsaP-overexpression on other T4aP proteins. Surprisingly, deletion of PilQ or the PilQ pilotin, PilF, did not prevent TsaP over-expression from increasing Congo-red dye uptake (**Figure 3D**). This indicates that PilQ is not downstream of TsaP in this signal cascade. Like TsaP, PilY1 over-expression in *trans* can trigger increased cyclic di-GMP production^12^. Deleting the *pilMNOP* genes blocks cyclic di-GMP induction upon PilY1 over-expression^12^. When TsaP was over-expressed in a *pilY1* deletion mutant, there was decreased Congo-red dye uptake, consistent with TsaP over-expression inducing a signal cascade that proceeds through PilY1 (**Figure 3D**). Likewise, TsaP over-expression in *pilN, pilO*, or *pilP* mutants resulted in less Congo red dye uptake (**Figure 3D**). These results are consistent with TsaP over-expression triggering a T4aP-related cyclic di-GMP signal transduction pathway.

## DISCUSSION

We determined the structure of PilQ alone and bound to TsaP, and found that TsaP over-expression leads to increased expression of cyclic di-GMP, upstream of PilY1. Our results resolve the identity of the spokes identified in Proteobacterial PilQ cryoEM densities and may reveal a new component of the surface-sensing signal cascade.

Where most secretins are bound by their cognate pilotin around the periphery of the secretin barrel adjacent to the secretin gate, we found PilQ^Pa^ is bound by TsaP. Unlike pilotins, TsaP is not essential for PilQ assembly or twitching motility^16,18,19^. Where and how the pilotin PilF interacts with PilQ remains an open question, although one possibility is that TsaP replaces PilF after insertion of PilQ into the outer membrane. We identified co-evolved residues that are consistent with the TsaP-binding region of PilQ interacting with the hydrophobic membrane-interacting interface. This hints at a model of PilQ membrane insertion in which secretin monomers initially fold onto themselves, with the TsaP-binding site occluding the hydrophobic membrane insertion site. Binding by pilotins to the TsaP-binding site might then promote the insertion of PilQ into the membrane, rationalizing the necessity of pilotins for secretin assembly and localization. If this were the case, after outer-membrane insertion of PilQ, PilF could be replaced by TsaP.

It seems likely that TsaP and PilQ are tightly associated, as the buried surface area of the TsaP-PilQ interaction is enormous (20468 Å^2^) and the complex is stable in detergent. It is nonetheless possible that some of the TsaP rings were removed during the purification, as only about half of the purified PilQ particles have TsaP associated with them. In this case, PilQ partially bound by TsaP might be expected, but only PilQ entirely bound by TsaP or without TsaP were identified in cryoEM 3D classes. Alternatively, there may be a population of free PilQ oligomers representing an intermediate state between PilQ-PilF and PilQ-TsaP complexes.

The structure of PilQ was determined in two distinct stoichiometries. Interestingly, only the tetradecameric form is bound by TsaP and had the unidentified secretin gate plug. The 2:1 PilQ:TsaP configuration necessitates binding of only even-numbered PilQ stoichiometries. It may be that TsaP recognizes or selects for specific secretin stoichiometries. Selecting particular secretin stoichiometries may improve the efficiency of T4aP function, consistent with the moderate twitching motility defect of the TsaP-deletion mutant.

In addition to influencing the quaternary structure of PilQ, TsaP binding may alter its tertiary structure. Comparing the different PilQ structures revealed a pivot point for secretin gate invagination adjacent to the spokes, consistent with TsaP binding influencing this process. Thus, a plausible role for TsaP may be in modifying secretin-gate invagination. The consequences of this are not clear, but the gate could then interact with the pilus initiation complex of minor pilins and PilY1 and/or the unidentified secretin gate plug.

TsaP could also be involved with larger-scale secretin movements. For example, PilQ adopts distinct open and closed states depending on whether the pilus fibre occupies the central channel the T4aP^19,20^. Compared to T4aP maps obtained by electron cryotomography in which the pilus fibre does not occupy the T4aP, the OM in the piliated state has moved 20 Å away from PilP and the peptidoglycan^19^. Although the secretin domain is embedded in the OM, the AMI_N and N0 domains of PilQ bind peptidoglycan and PilP, respectively^41^. This suggests that PilQ must extend and contract 20 Å between the piliated and non-piliated states. As TsaP binds to both the peptidoglycan and secretin domain of PilQ, TsaP binding could modulate secretin opening. For example, TsaP could act as an elastic band between the peptidoglycan and outer-membrane components of PilQ, keeping PilQ contracted and the T4aP closed in the absence of an assembled pilus. The predicted disordered residues between the LysM and CI/CII domains could afford TsaP the elasticity required for this action.

Alternatively, instead of modulating PilQ structure, TsaP may sense PilQ conformations and relay this information as a part of a signal cascade. However, while TsaP over-expression triggers the cyclic di-GMP signal cascade, it does so independently of PilQ. This is curious as PilQ provides the only known physical contact between TsaP and PilN, PilO, and PilP – which are important for inducing cyclic di-GMP production. It may be that there is a PilQ-independent signal transduction pathway from TsaP to PilY1 and PilNOP. Alternatively, over-expressing TsaP may force a signalling event that is typically dependent on PilQ. As TsaP has a peptidoglycan-binding LysM domain, overexpression might place strain on the peptidoglycan that resembles what happens during surface interactions, leading to cyclic di-GMP induction. In the absence of a known trigger for the signalling cascade upstream of TsaP, it is not clear what role TsaP plays in the surface-sensing signal cascade under native expression conditions. Nonetheless, knowing that TsaP over-expression can trigger the cyclic di-GMP signal cascade upstream of PilY1 will be useful in identifying other key players in the surface sensing signal cascade.

## METHODS

### Cloning and mutagenesis

TsaP from *P. aeruginosa* PAO1 was PCR amplified from genomic DNA using primers TsaP_fwd and TsaP_rev (**Supplementary Table 3**), digested with KpnI and XbaI, and cloned into pBADGr to create pBADGr:*tsaP* for arabinose inducible expression of TsaP. Plasmid sequences were verified by TCAG sequencing facilities (The Hospital for Sick Children, Canada).

### TsaP Deletion

A gene deletion allele was ordered from BioBasic and subcloned into pEX18Gm at the BamHI site producing pEX18Gm_*tsaP*. Clones were confirmed by digest and DNA sequencing (TCAG, The Hospital for Sick Children). The pEX18Gm_*tsaP* construct was transformed into chemically competent SM10 *E. coli* cells, colonies containing pEX18Gm_*tsaP* were mated with mPAO1 using a method described elsewhere^42^. Briefly, after mating *P. aeruginosa* was selected for by growth on LB agar plates containing 25 μg/ml triclosan (Sigma) followed by selection of the merodiploid strains on VBMM agar containing 30 μg/ml gentamicin. Counter selection to resolve the merodiploids was performed on 1.5 % agar plates with 10 g/L tryptone, 5 g/L yeast extract, and 15% (wt/vol) sucrose. The deletion was verified by the absence of a PCR product using primers TsaP_fwd and TsaP_rev.

### Phenotypic characterizations

*P. aeruginosa* PAO1 with and without deletion mutations of *tsaP, pilN, pilO, pilP, pilQ, pilY1* were electroporated with pBADGr or pBADGr:tsaP. *P. aeruginosa* PAO1 with and without deletion mutations of *tsaP* were also electroporated with pBADGr:tsaP.

Twitching assays were performed as described previously^43,44^ except that assays were performed in 150 mm by 15 mm petri polystyrene dishes (Fisher Scientific) with 30 µg/ml gentamicin for 18 h at 37 °C. After this incubation, the agar was then carefully discarded and the adherent bacteria were stained with 1 % (w/v) crystal violet dye, followed by washing with deionized water to remove unbound dye. Twitching zone areas were measured using ImageJ software^45^. Twitching motility assays were performed in six replicates.

For Congo-red dye binding assays, cells were grown in 0.5 mL of lysogeny broth (LB) with 30 µg/ml gentamicin for 16 h at 37 °C while shaking. Cultures were normalized to an OD at 600 nm of 0.1 and then 3 μl of culture was spotted onto Congo red plates (1% (wt/vol) Tryptone, 1% (wt/vol) agar, 0.004% (wt/vol) congo red, and 0.0015% (wt/vol) Coomassie Brilliant Blue R250, and either 0, 0.01, or 0.1% L-arabinose) with the appropriate antibiotic, and grown for 24-36 h at 37 °C.

Extracellular proteases were assayed using skim milk plates (1% wt/vol agar, 1% wt/vol skim milk powder, and either 0, 0.01, or 0.1% L-arabinose) with the appropriate antibiotic, using the same normalized cultures used for the Congo red binding assays. 3 μl of culture was spotted on a plate and incubated for 16 hrs at 37 °C. All plates were scanned using an Epson Perfection 4990 scanner and Epson Scan software.

For luciferase expression, cells previously electroporated with pBADGr, pBADGr::tsaP and pBADGr::tsaP mutant constructs were electroporated with pMS402:PcdrA. Cells were grown in 0.5 mL of lysogeny broth (LB) with 30 µg/ml gentamicin and 150 µg/ml kanamycin for 16 h at 37 °C while shaking. Cultures were normalized to OD600 of 0.1 and diluted 10x into fresh LB broth containing 30 μg/ml gentamicin, 105 μg/ml kanamycin and either 0, 0.01% or 0.1% L-arabinose in a 384 well plate (Corning, 3821). All samples were done in triplicate. Growth and luciferase activity were monitored over a 16-hour period in a Synergy Neo2 plate reader (BioTek Instruments). Data was analyzed and plotted using Prism 8 (Graph Pad).

### Plotting the residues at the packing unit interface

CMview^46^ was used to identify residues that are near one another. The contact type was set to ‘ALL’ (i.e. every atom available in the structure) and the distance cut-off was set to 8 Å. To identify residues that are near one another in PilQ monomers, the CI domain of TsaP (residues 126-211), or the CII domain of TsaP (residues 235-324), these structure segments were loaded separately. To identify intra-molecular CI to CII residues that are near each other in TsaP, full-length TsaP was loaded. To identify inter-molecular residues that are near one another, two adjacent PilQ protomers were loaded together, or the CI domain and CII domains from two adjacent protomers were loaded together.

### Identifying evolutionarily-coupled residues

The TsaP and PilQ^Pa^ amino acid sequences were analyzed using the EVcouplings webserver^26^ with default parameters. Homologs were identified, aligned, and analyzed – in total, 2642 and 11302 for TsaP and PilQ, respectively. There was poor alignment quality in the lip subdomain of PilQ (residues 481-531) so few evolutionarily-coupled residue pairs were discovered in the lip subdomain. Only residue pairs with a probability score greater than 0.845 or 0.985 were included in subsequent analysis, for TsaP and PilQ, respectively. To better understand the significance of these evolutionarily coupled residues, they were compared to the residues that are near one another identified in CMview.

### CryoEM analysis

As previously described for this dataset^18^, cryoEM data was collected as movies (30 frames over 15 sec) with a FEI Tecnai F20 electron microscope operating at 200 kV and equipped with a Gatan K2 Summit direct detector camera. The exposure rate was 5 electrons/pixel/s with a calibrated pixel size of 1.45 Å/pixel. Previously, movie frames had been aligned with alignframes_lmbfgs^47^ and then individual movie frames were discarded for data storage purposes; this prevented exposure weighting and individual particle motion correction herein. Further image processing here was performed in cryoSPARC v2.8.0^24^ (**Supplementary Table 1**). CTF parameters were estimated from the average of aligned frames with CTFFIND4^48^ within cryoSPARC and individual particle motion correction and exposure weighting was done with an implementation of alignparts_lmbfgs^47^ within cryoSPARC. Initial 2D class averages were generated with manually selected particle images; these class averages were then used to select more particle images. Particle images were extracted in 256×256 pixel boxes then subjected to 2D classification. For the purposes of 3D classification, particle images contributing to 2D classes without high-resolution features were removed. *Ab initio* reconstruction was performed using two classes, revealing barrels and spoked-barrels. The particles contributing to the barrels without spokes were used for an additional round of *ab initio* reconstruction with two classes, revealing an additional class with wider barrels. The spoked barrels, the barrels without spokes, and the wider barrels without spokes were then used as initial models for heterogeneous refinement; particles from 3D classes that did not converge at this stage were removed. Particles from distinct 3D classes were then subjected to non-uniform refinement^49^. The resulting fourier shell correlations for the 4.3 and 4.4 Å maps are unusual as there is a flat region between ∼7 Å and ∼5 Å. This likely arises from the diffuse density from the detergent layer and the N-terminal domains disordered relative to the secretin of PilQ. However, the clear features from the beta-barrels support the resolution estimates for the maps

A molecular model could be built using Coot^50^ into the spoked barrel map using the structures of FlgT and GspD^25,32^ as guides for TsaP and PilQ, respectively. Molecular models for the maps of barrels without spokes were built by fitting the PilQ protomers built into the spoked barrel map in Coot^50^. These models were refined against the maps in Phenix-Refine^51^ with reference model restraints to the spoked barrel PilQ structures. While side-chains were used to ensure the models made biological sense during model building, they were ultimately removed as the map quality did not justify them being modeled.

## Supporting information

Animation 1

## Acknowledgements

This work was supported by a grant from the Canadian Institutes for Health Research (CIHR) MOP 93585 to L.L.B. and P.L.H. M.M. was supported by a CIHR Doctoral Studentship and Ontario Graduate Scholarship during these studies. P.L.H. and J.L.R. are recipients of Canada Research Chairs. The graphics programs used in this study were accessed using SBGrid.

