## Supplemental Tables & Figures for "CryoEM map of *Pseudomonas aeruginosa* PilQ enables structural characterization of TsaP"

**Supplementary Table 1. CryoEM data processing, atomic model statistics, and map/model depositions.**

| **Image processing** | **All other samples** | | |
|---|---|---|---|
| Motion correction software | cryoSPARC v2 | | |
| CTF estimation software | CTFFIND4 | | |
| Particle selection software | cryoSPARC v2 | | |
| Classification and refinement software | cryoSPARC v2 | | |
| Micrographs | 2351 | | |
| Selected particles | 324516 | | |
| **EM maps** | **$C_{14}$ Barrel $C_7$ spokes** | **$C_{14}$ Barrel** | **$C_{15}$ Barrel** |
| Particle images contributing to maps | 55006 | 54040 | 16914 |
| Applied symmetry | $C_7$ | $C_{14}$ | $C_{15}$ |
| Applied B-factor ($Å^2$) | -98 | -239 | -671 |
| Global resolution (FSC = 0.143, Å) | 4.3 | 4.4 | 6.9 |
| **Model building** | **$C_{14}$ Barrel $C_7$ spokes** | **$C_{14}$ Barrel** | **$C_{15}$ Barrel** |
| Modeling software | Coot , ISOLDE, Phenix, Rosetta | Coot , ISOLDE, Phenix, Rosetta | Coot , ISOLDE, Phenix, Rosetta |
| Atoms | 30198 | 22484 | 22425 |
| RMSD | | | |
| Bond length (Å) | 0.007 | 0.009 | 0.012 |
| Bond angle (°) | 1.53 | 1.85 | 1.75 |
| Ramachandaran | | | |
| Favoured (%) | 95 | 95 | 95 |
| Allowed | 99 | 99 | 99 |
| Clashscore | 17 | 26 | 28 |
| Av. B-factor ($Å^2$) | 204 | 275 | 752 |
| *MolProbity* score[‡] | 2.1 | 2.3 | 2.3 |
| $CC_{mask}$ | 0.7 | 0.8 | 0.7 |
| **Deposited maps and coordinate files** | | | |
| EMDB code | EMD-21152 | EMD-21153 | EMD-21154 |
| PDB code | 6VE2 | 6VE3 | 6VE4 |

[‡], As calculated using *MolProbity*[45] implemented in *PHENIX*

**Supplementary Table 2. Summary of ten most abundant proteins as identified by trypsin digest followed by mass spectrometry.**

| Protein name | Number of unique peptides | Total peptide count | % coverage |
|---|---|---|---|
| PrpL | 56 | 327 | 75 |
| TsaP | 39 | 88 | 74 |
| PilQ | 16 | 21 | 70 |
| PA5122 | 13 | 34 | 75 |
| ArcA | 9 | 11 | 16 |
| PA4078 | 8 | 10 | 2 |
| PA1053 | 5 | 18 | 42 |
| PA5333 | 5 | 12 | 23 |
| RplN | 5 | 11 | 23 |
| PA0359 | 5 | 9 | 14 |

**Supplementary Table 3: Primers, strains, and plasmids used in this study**

| **Primers** | **Sequence** | **Reference** |
|---|---|---|
| TsaP_fwd | TATAATGGTACCCATGAGGAAATCACTAGTCGCCCTTC | This Study |
| TsaP_rev | TATATATCTAGATCAGGGATTCTGCACCC | This Study |
| **Strains** | **Description** | **Reference** |
| *E. coli* TOP10 | $F^-$, mcrA Δ(mrr-hsdRMS-mcrBC) φ80lacZΔM15 ΔlacX74 nupG recA1 araD139 Δ(ara-leu)7697 galE15 galK16 rpsL($Str^R$) endA1 $\lambda^-$ | Invitrogen |
| SM10 | *thi-1 thr leu tonA lacY supE recA::RP4-2-Tc::Mu* ($Km^R$) | Invitrogen |
| mPAO1 | WT | [46] |
| *P. aeruginosa* PAO1 Δ*tsaP* | Deletion of *tsaP* | This Study |
| *P. aeruginosa* PAO1 Δ*pilN* | Transposon insertion mutant in *pilN* at 271 | [46] |
| *P. aeruginosa* PAO1 Δ*pilO* | Transposon insertion mutant in *pilO* at 226 | [46] |
| *P. aeruginosa* PAO1 Δ*pilP* | Transposon insertion mutant in *pilP* at 254 | [46] |
| *P. aeruginosa* PAO1 Δ*pilQ* | FRT scar insertion mutant in *pilO* at 571 | [47] |
| *P. aeruginosa* PAO1 Δ*pilF* | Transposon insertion mutant in *pilF* at residue 633 | [46] |
| *P. aeruginosa* PAO1 Δ*pilY1* | Transposon insertion mutant in *pilY1* at 1407 | [46] |
| *P. aeruginosa* PAO1 Δ*pilA* | Transposon insertion mutant in *pilA* | [46] |
| **Plasmids** | **Description** | **Reference** |
| pBADGr | pBAD plasmid with gentamicin resistance | [48] |
| pBADGr:*tsaP* | TsaP from *P. aeruginosa* for expression in *P. aeruginosa* | This Study |
| pMS402:P*cdrA* | Lux genes with the cyclic di-GMP inducible *cdrA* promoter. | [8] |
| pEX18Gm_*tsaP* | Deletion construct for mPAO1 Δ*tsaP* | This Study |

```
  1 MNSGLSRLGIALLAAMFAPALLAADLEKLDVAALPGDRVELKLQFDEPVAAPRGYTIEQP 60
 61 ARIALDLPGVQNKLGTKNRELSVGNTRSVTVVEAKDRTRLIINLTALSSYTTRVEGNNLF 120
121 VVVGNSPAGASVASAAPVKASPAPASYAQPIKPKPYVPAGRAIRNIDFQRGEKGEGNVVI 180
181 DLSDPTLSPDIQEQGGKIRLDFAKTQLPDALRVRLDVKDFATPVQFVNASAQSDRTSITI 240
241 EPSGLYDYLVYQTDNRLTVSIKPMTTEDAERRKKDNFAYTGEKLSLNFQDIDVRSVLQLI 300
301 ADFTDLNLVASDTVQGNITLRLQNVPWDQALDLVLKTKGLDKRKLGNVLLVAPADEIAAR 360
361 ERQELEAQKQIAELAPLRRELIQVNYAKAADIAKLFQSVTSDGGQEGKEGGRGSITVDDR 420
421 TNSIIAYQPQERLDELRRIVSQLDIPVRQVMIEARIVEANVGYDKSLGVRWGGAYHKGNW 480
481 SGYGKDGNIGIKDEDGMNCGPIAGSCTFPTTGTSKSPSPFVDLGAKDATSGIGIGFITDN 540
541 IILDLQLSAMEKTGNGEIVSQPKVVTSDKETAKILKGSEVPYQEASSSGATSTSFKEAAL 600
601 SLEVTPQITPDNRIIVEVKVTKDAPDYQNMLNGVPPINKNEVNAKILVNDGETIVIGGVF 660
661 SNEQSKSVEKVPFLGELPYLGRLFRRDTVTDRKNELLVFLTPRIMNNQAIAIGR       714
```

**Supplementary Figure 1.** Mass spectrometry peptide coverage of PilQ. Peptides present in the mass spectrometry data are shown in yellow. The hydrolyzed secretion signal sequence is underlined.

1 MRKSLVALLLLAASGLAQAQVDLREGHPDRYTVVRGDTLWDISGKFLRQPWKWPELWHAN 60
61 PQIQNPHLIYPGDTLSLVYVDGQPRLVLNRGESRGTIKLSPKIRSTPIAEAIPTIPLDKI 120
121 NSFLLANRIVDDEKTFTSAPYIVAGNAERIVSGTGDRIYARGKFADGQPAYGIFRQGKVY 180
181 IDPKTKEVLGINADDIGGGEVVATEGDVATLALTRTTQEVRLGDRLFPTEERAVNSTFMP 240
241 GEPSREVKGEIIDVPRGVTQIGQFDVVTLNRGQRDGLAEGNVLAIYKVGETVRDRVTGES 300
301 VKIPDERAGLLMVFRTYKKLSYALVLMASRPLSVTDRVQNP 341

**Supplementary Figure 2.** Mass spectrometry peptide coverage of TsaP. Peptides present in the mass spectrometry data are shown in yellow. The hydrolyzed secretion signal sequence is underlined.

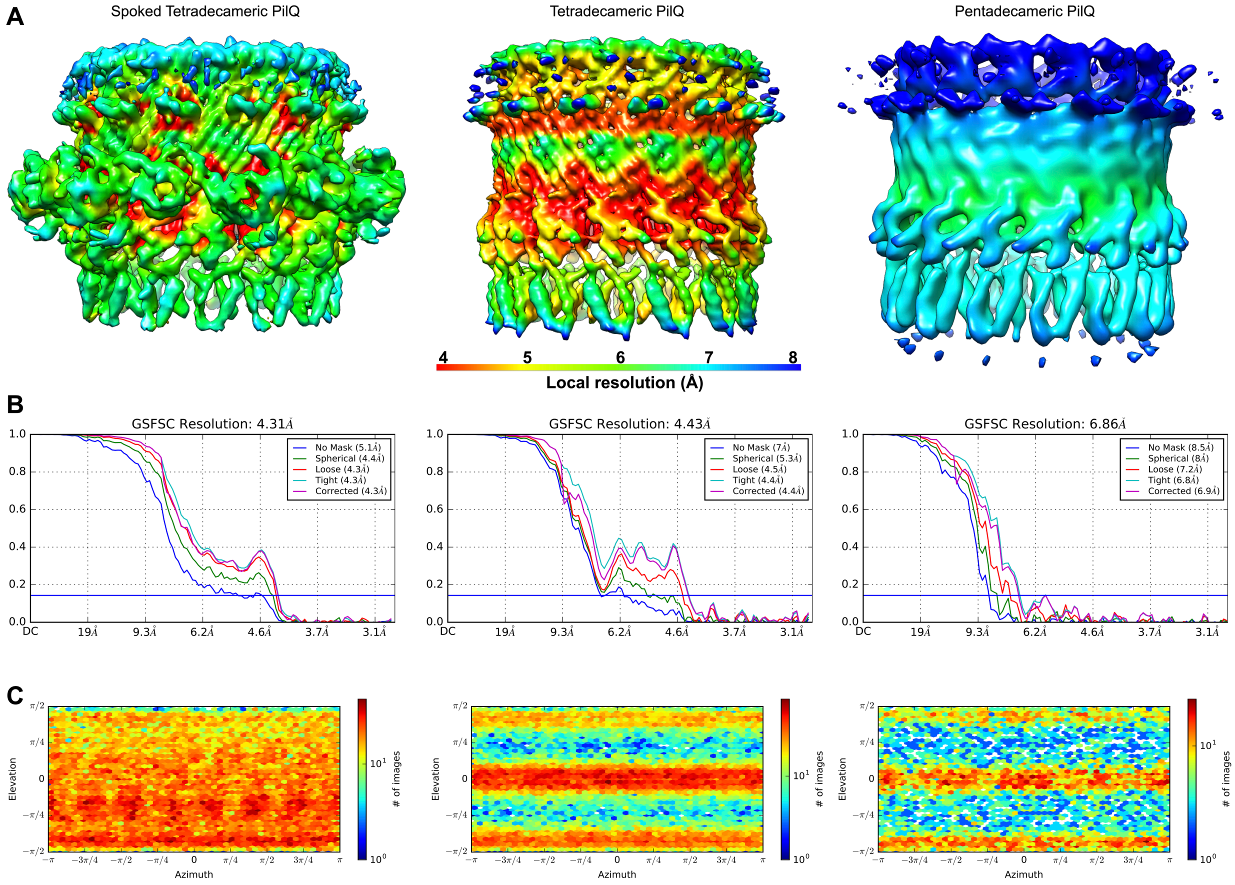


**Supplementary Figure 3.** Local resolution (**A**), Fourier shell correlation (FSC) curves (**B**), and particle distribution (**C**).
